# Oral OM-89 in combination with antibiotics prevents recurrent infection in a mouse model of urinary tract infection

**DOI:** 10.1101/2025.04.04.647233

**Authors:** Tracy Canton, Méline Durand, Edouard Baulier, Christian Pasquali, Ruxandra Calin, Molly A Ingersoll, Matthieu Rousseau

**Affiliations:** Department of Immunology, Institut Pasteur, Paris France; Mucosal Inflammation and Immunity Team, Université Paris Cité, CNRS, Inserm, Institut Cochin and Department of Immunology, Institut Pasteur, Paris France; OM Pharma SA, Preclinical Research Department, 1217 Meyrin, Geneva, Switzerland; Sorbonne University Tenon Hospital, Paris, France

## Abstract

Urinary tract infection (UTI) is a very common infection. Approximately 25% of all women will experience recurrent infection, defined by 2 UTI in 6 months or 3 UTI in a 12 month period. Recurrent infection is thought to be due in part to the development of local bacterial reservoirs in the bladder after an infection. These reservoirs are undetected by the host immune system and may re-emerge after resolution of the primary infection. Frequently, recurrent UTI require repeated antibiotic use, leading to the development of resistance, and negatively impacting the quality of life of the patient. Given the rise in multidrug resistant uropathogens, treatments that do not rely upon antibiotics are urgently needed for this patient population. We assessed the capacity of a lyophilized *E. coli* extract, OM-89®, as a preventive measure to reduce the incidence of spontaneous recurrence in a mouse cystitis model. Administration of OM-89 *per os* after the initiation of an acute UTI significantly reduced the number of spontaneous recurrent UTI in female mice over a 1 month follow-up period only when the animals were also treated with antibiotics. Protection against recurrent UTI arose quickly and without changes in any adaptive immune response parameter measured, ruling out a role for OM-89 in augmenting adaptive immunity to protect against spontaneous recurrent UTI in the first month of use.

## Introduction

UTI occurs in approximately 50% of all women during their lifetime, and recurrent UTI arises in nearly 50% of those experiencing a first infection (Yang et al., 2022). While definitions vary from country to country, generally, recurrent UTI is clinically defined as having two infections in a 6 month period or three infections in a 12 month period (Kranz et al., 2024). Mechanisms of recurrence are poorly understood and are likely multi-factorial. Evidence from bladder biopsies of recurrent UTI patients supports that uropathogens can form reservoirs in human tissue (De Nisco et al., 2019; Gadhvi et al., 2024; Rosen et al., 2007). In mice, uropathogens, such as uropathogenic *E coli* (UPEC) and *Klebsiella pneumoniae* form quiescent reservoirs in the intermediate layers of the urothelium, where they are protected from soluble and cellular host responses and antibiotic treatment, and spontaneously re-enter the acute pathogenic cycle at varying time points after a first infection, leading to a second UTI (Amoura et al., 2024; Anderson et al., 2004; Mysorekar and Hultgren, 2006; Rosen et al., 2008; Schilling et al., 2002). Thus, uropathogen reservoirs may contribute to recurrent UTI, however, the kinetics and signals that induce bacterial reactivation and infection are not known.

Recurrence may also arise following transition of commensal bacteria, such as *E. coli*, from the gut to the bladder (Yamamoto et al., 1997). Longitudinal multiomics analyses in women with recurrent UTI revealed that while their gut microbiomes are distinct from women without UTI, relative abundance of *E. coli* was not different between the groups (Worby et al., 2022). Despite the lack of difference in abundance, this study does not rule out that gut dysbiosis, potentially mediated by long-term or repeated antibiotic therapy, increases susceptibility to UTI caused by commensal *E. coli* (Worby et al., 2022).

Finally, recurrence may also arise due to inability of the mucosal immune response to resolve the infection (Lacerda Mariano et al., 2020; Mora-Bau et al., 2015; Rousseau et al., 2023). Development of immune memory, after infection or by immunomodulation, such as vaccination, protects a host from having the same infection more than once. However, sterilizing immune memory often fails to develop, particularly at mucosal sites, for reasons that are incompletely understood, leading to recurrent infection. This failure can lead to recurrent infection by pathogens that are phylogenetically close or identical to the original bacterial strain, including in UTI (Ikaheimo et al., 1996). Identifying a means to prevent recurrence would contribute to reducing the negative health and economic impact of this disease and be expected to reduce clinical reliance on antibiotics.

OM-89, sold under the brand name Uro-Vaxom®, is a lyophilizate produced by complex chemical lysis manufacturing. The resultant product is in an extract of proprietary bacterial molecules derived from degraded bacterial lysates of 18 *E. coli* strains. The product is used in a capsule formulation developed to prevent recurrent UTI in humans. A meta-analysis of five clinical studies demonstrated that OM-89 is safe, well-tolerated, and significantly reduces the number of recurrent UTI experienced by women when taken daily (Bauer et al., 2002). A randomized, double-blind, placebo-controlled trial of 453 women with a history of recurrent UTI found that individuals treated with OM-89 had a 34% reduction in infection, which was statistically significantly reduced compared to the placebo group over a 12 month follow-up period (Bauer et al., 2005). In the study design, patients were treated with OM-89 daily for 90 days, stopped treatment for 90 days, and resumed treatment on the first 10 days of the month for an additional 3 months. A more recent meta-analysis evaluated nine studies using OM-89, including seven randomized controlled trials, a retrospective cohort study, and a cross over trial (Prattley et al., 2020). Eight studies included a comparator group of either control (one 50 mg capsule of nitrofurantoin, once per day for 3 months) or placebo (similar capsule) and followed the design of 3 month treatment with or without a 10-day/month booster between months 6 to 9 (Prattley et al., 2020). Globally, OM-89 treatment reduced UTI between 52.6% and 87.5% compared to 50% in the placebo group. Overall, OM-89 showed a significantly improved odds ratio in the short term (<6 months), 0.29 (95% CI 0.10–0.87). The updated 2025 EAU guidelines recommend further study of promising agents, such as OM-89 in larger, high quality randomized clinical trials, using standardized definitions of recurrent cystitis and other outcome measures (*EAU Guidelines. Edn. presented at the EAU Annual Congress, Madrid 2025. ISBN 978-94-92671-29-5*.).th

Thus, although OM-89 is considered to be immunomodulatory, how it reduces recurrent UTI is unknown. To improve upon its efficacy, and potentially refine its mechanism of action, an animal model reflecting human responses is needed. Our objective was to establish a mouse model of OM-89 treatment to determine whether oral administration of this drug could prevent spontaneous recurrent UTI in mice, as a first step to understanding its mechanism of action. We found OM-89 treatment prevented spontaneous recurrent UTI, however, only in the context of antibiotic treatment. OM-89 treatment did not induce detectable changes in innate immune responses shortly after infection or in antibody production or adaptive immune cell infiltration, development, or accumulation, ruling out that OM-89 prevents spontaneous or intentional recurrent UTI by strengthening the adaptive immune response to infection.

## Results

### OM-89, in combination with antibiotic treatment, reduces spontaneous recurrent UTI

To test whether OM-89 could prevent spontaneous recurrent UTI in a laboratory setting, we infected female C57BL/6 mice with 10^7^ colony forming units (CFU) of uropathogenic *E. coli* (UPEC) strain UTI89-RFP-kanR (Mora-Bau et al., 2015; Zychlinsky Scharff et al., 2017). At 24 hours post-infection (PI), cohorts of mice were left untreated, or treated with OM-89, trimethoprim/sulfamethoxypyridazine (TMP-SMX) in the drinking water for 5 days, a single dose of fosfomycin per os, TMP-SMX and OM-89, or fosfomycin and OM-89 (TMP-SMX/OM-89 or fos/OM-89, respectively) (**Figure 1A**). Our primary objective, to monitor resolution of infection and spontaneous re-infection over time, was achieved by urine collection twice weekly with qualitative monitoring for the presence of UPEC. OM-89-treated mice resolved acute UTI with the same kinetic as untreated animals (**Figure 1B**). Surprisingly, we observed a 67% reduction in the total number of recurrence events between TMP-SMX-treated and TMP-SMX/OM-89 treated mice, and a 100% reduction between fosfomycin-treated and fos/OM-89-treated mice during a 28 day follow-up period (**Figure 1B-C**). By contrast, no differences in the number of recurrent events were observed in mice treated only with OM-89 compared to their matched, untreated control group.

**Figure 1:**
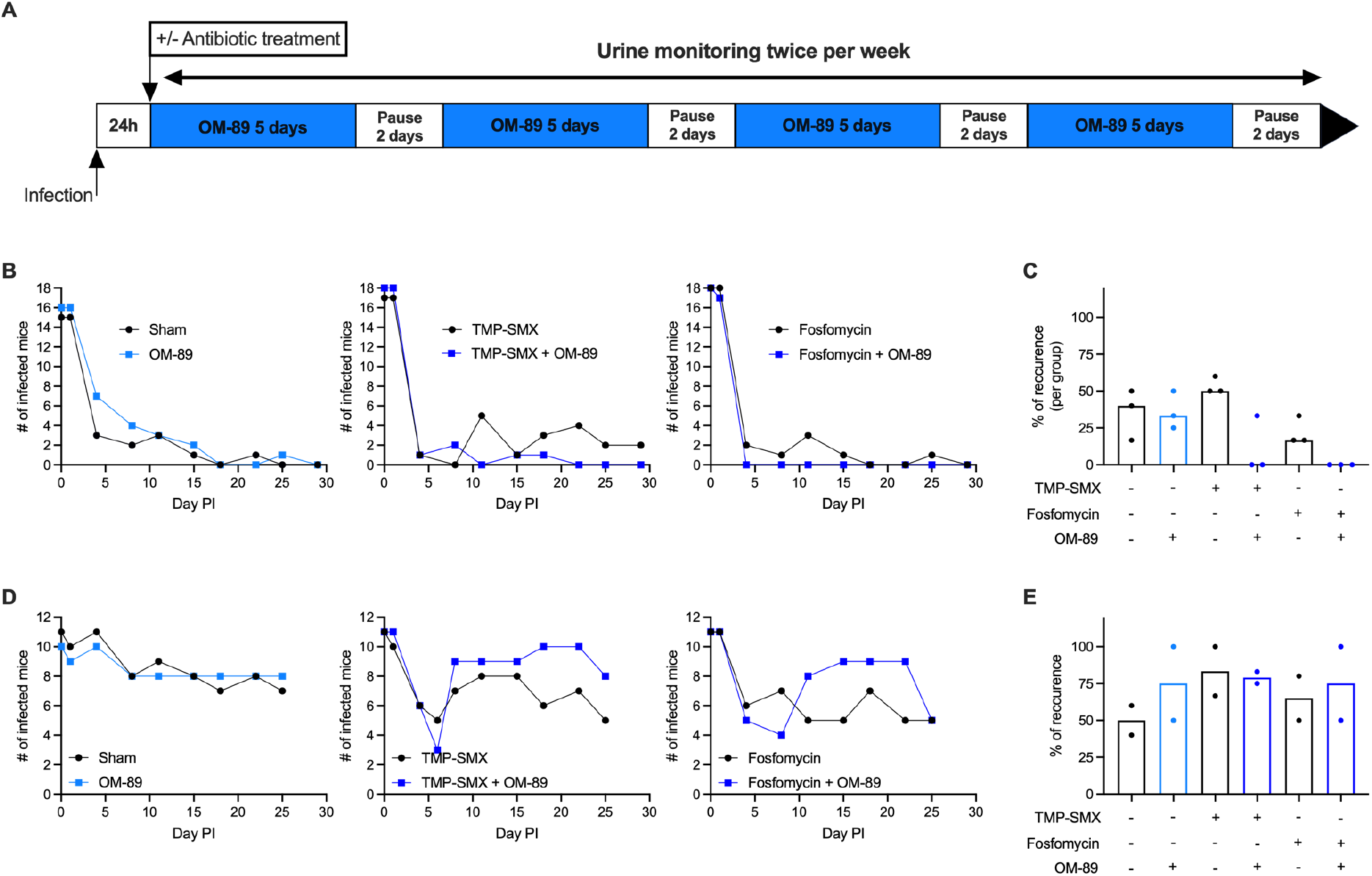
OM-89, in combination with antibiotic therapy, prevents spontaneous recurrent UTI in female mice. (A) Experimental design scheme shows treatment and sample collection schedule. (**B-E**) Mice were infected with 10^7^ CFU of UPEC strain UTI89-RFP-KanR and treated as shown in (**A**). Graphs show the number of infected animals and proportion of recurrent infection events in (**B-C**) female mice or (**D-E**) male mice. Data in (**B**) are pooled from 3 independent experiments, n=6/experiment, data in (**D**) are pooled from 2 independent experiments, n=6/experiment. Each dot in (**C, E**) represent one experiment.

We performed the same experiment in male mice, and as expected from our previous work, untreated male mice remained persistently infected for 25 days (**Figure 1D**). Interestingly, TMP-SMX, TMP-SMX/OM-89, and fos/OM-89 treated mice all rebounded in the three days following antibiotic treatment, illustrating the challenges of treating male UTI (**Figure 1D**). OM-89 did not improve resolution kinetics in any group and potentially impeded bacterial clearance in male mice treated with antibiotics, as more mice had positive urine cultures in the TMP-SMX/OM-89 and fos/OM-89 treatment groups compared to antibiotics alone (**Figure 1D**). The number of recurrent infections in male mice was not different in any experimental group (**Figure 1E**). As OM-89, with or without antibiotics, had no impact on male mice, and male mice remained chronically infected, we did not perform further experimentation in male mice.

### OM-89 treatment does not alter antibody concentrations in serum or bladders

We hypothesized that OM-89 and antibiotic therapy increases antibody production to reduce recurrent UTI. We focused on fos/OM-89 treatment as this combination completely prevented recurrent events in the follow-up period (**Figure 1B**). We noted that when mice had recurrent events, these began at day 11 PI (**Figure 1B**). Thus, mice were infected and treated, and blood was collected at day 0, 5, and 11 PI, at which point the animals were sacrificed (**Figure 2A**). We measured the lambda (**Figure 2**) and kappa (**Supplementary figure 1**) light chains of IgA, IgG1, IgG2a, IgG2b, IgG3, and IgM in the serum and bladder using multiplexed analysis. We observed no significant differences between the fosfomycin group and the fos/OM-89 group at any timepoint in the serum (**Figure 2B and Supplementary figure 1**) or in the bladder at day 11 (**Figure 2C and Supplementary figure 1**).

**Figure 2:**
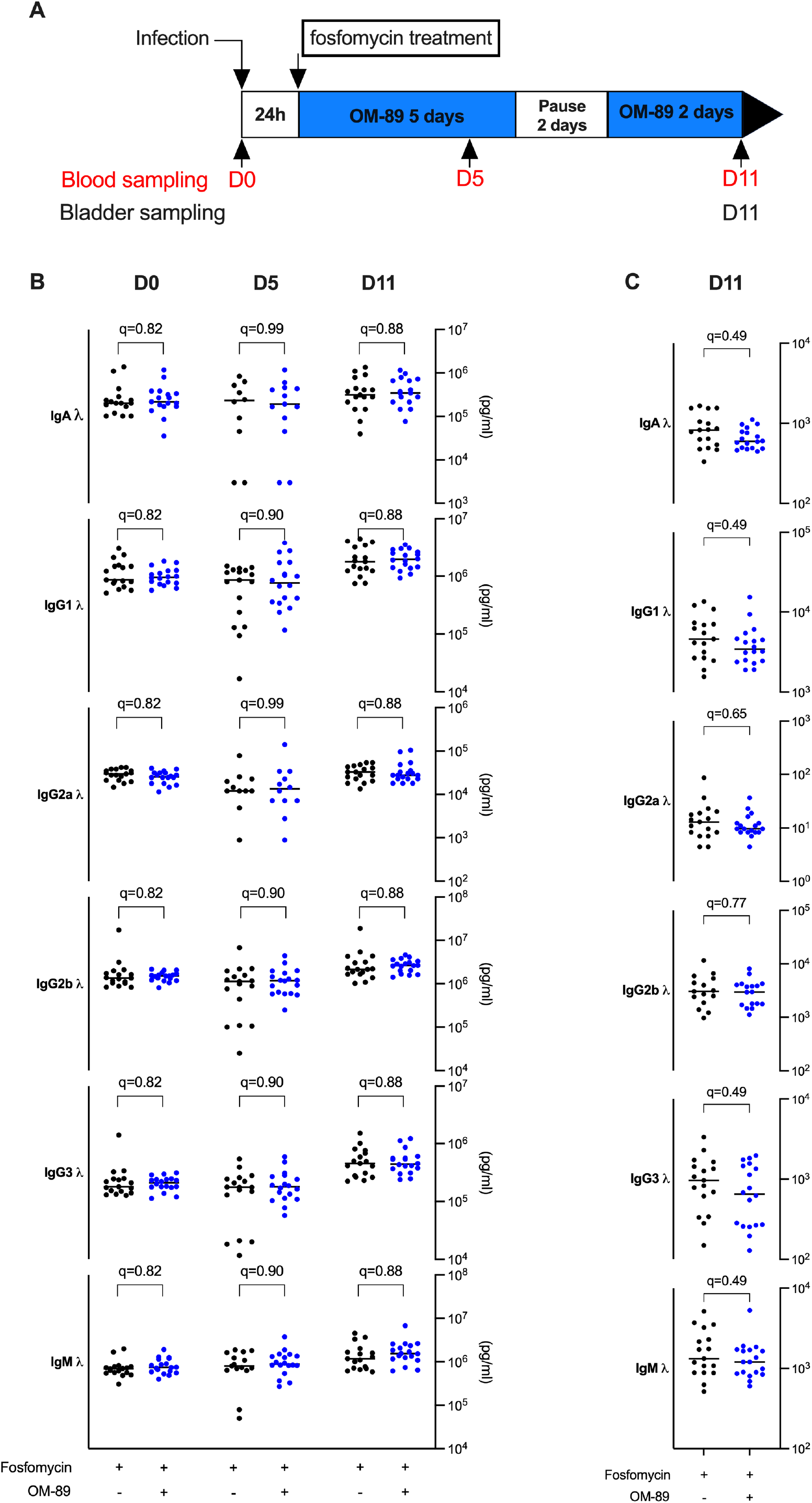
OM-89 does not alter antibody levels in serum or bladder. (**A**) Female mice were infected and then treated with fosfomycin or fos/OM-89 according to the experimental scheme. Graphs show antibody concentrations in (**B**) serum on day 0, 5, and 11 PI and (**C**) bladder homogenates on day 11 PI. Data are pooled from 3 independent experiments; n=6 mice/experiment. Each dot is a mouse and lines are medians. Nonparametric Mann-Whitney tests comparing fosfomycin treatment to fos/OM-89 treatment for each immunoglobulin concentration were performed. P values were corrected for multiple comparisons using the FDR method and all q values are shown.

### OM-89 treatment does not alter immune cell populations in the blood

Having ruled out an impact on antibody production, we hypothesized that OM-89 treatment, in combination with antibiotics, may impact circulating immune cells, which would then be available to respond to a recurrence event. Mice were infected, treated and sampled as in **Figure 2A**. As expected, no significant differences were observed among the groups at day 0 (naïve mice) (**Figure 3**). We also observed no significant differences in immune cell numbers between the fosfomycin group and the fos/OM-89 group at day 5 or 11 PI in the serum (**Figure 3**).

**Figure 3:**
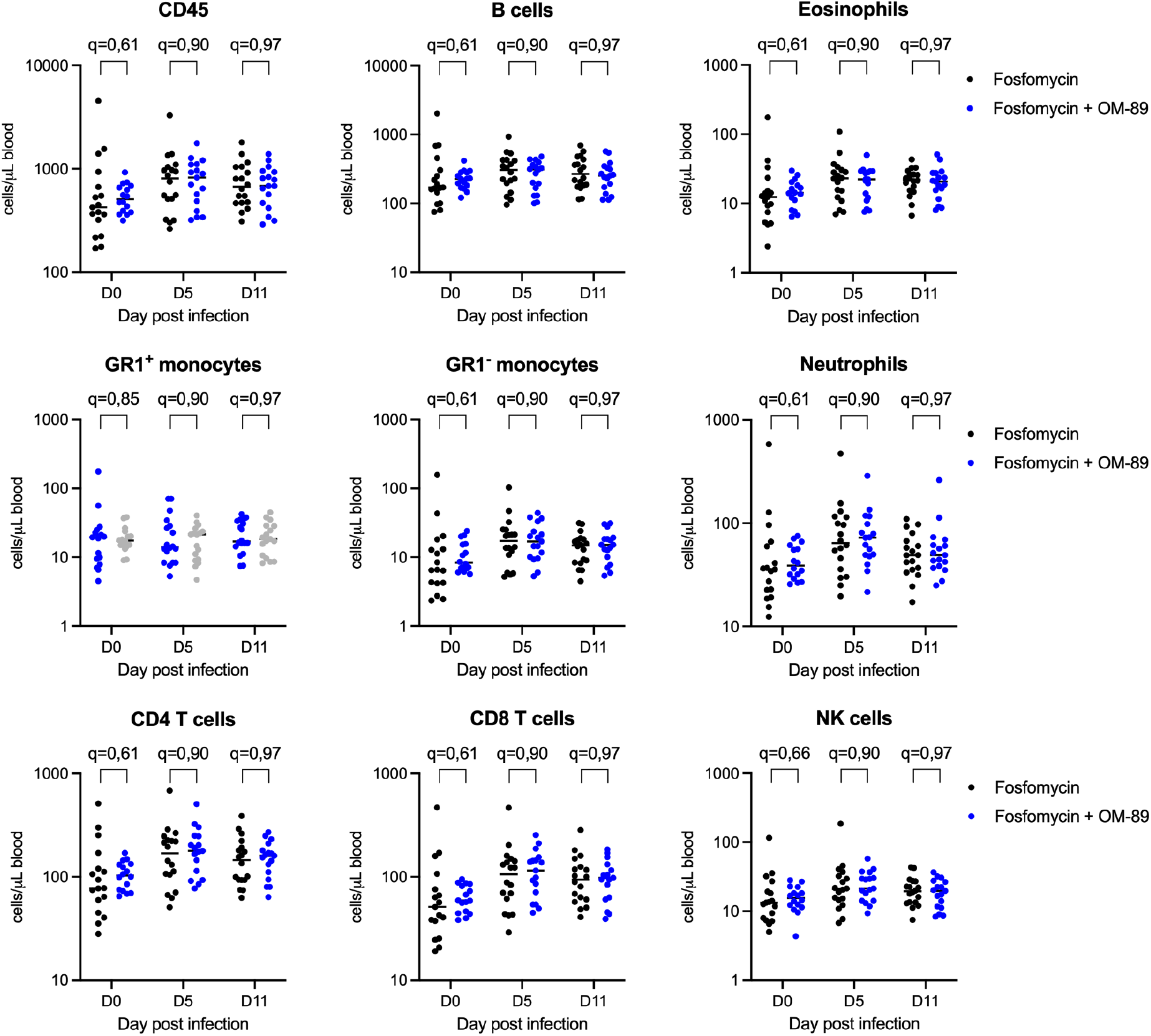
OM-89 treatment does not alter circulating immune cell numbers in the blood. Female mice were infected, treated, and sampled as in Figure 2A. Graphs depict the concentration of indicated immune cells per/μl of blood. Data are pooled from 3 independent experiments; n=6 mice/experiment. Each dot is a mouse and lines are medians. Nonparametric Mann-Whitney tests comparing fosfomycin treatment to fos/OM-89 treatment for each immune cell were performed. P values were corrected for multiple comparisons using the FDR method and all q values are shown.

### OM-89 treatment does not change immune cell infiltration into the bladder

As we did not observe a systemic impact on circulating immune cell numbers, we hypothesized that fos/OM-89 treatment acted locally in the bladder mucosa, potentially increasing immune cell infiltration. We quantified myeloid and lymphoid cell infiltration at 11 days PI and observed no significant differences in immune cell infiltration in any subset between the fosfomycin group and the fos/OM-89 group at 11 PI in the bladder (**Figure 4**).

**Figure 4:**
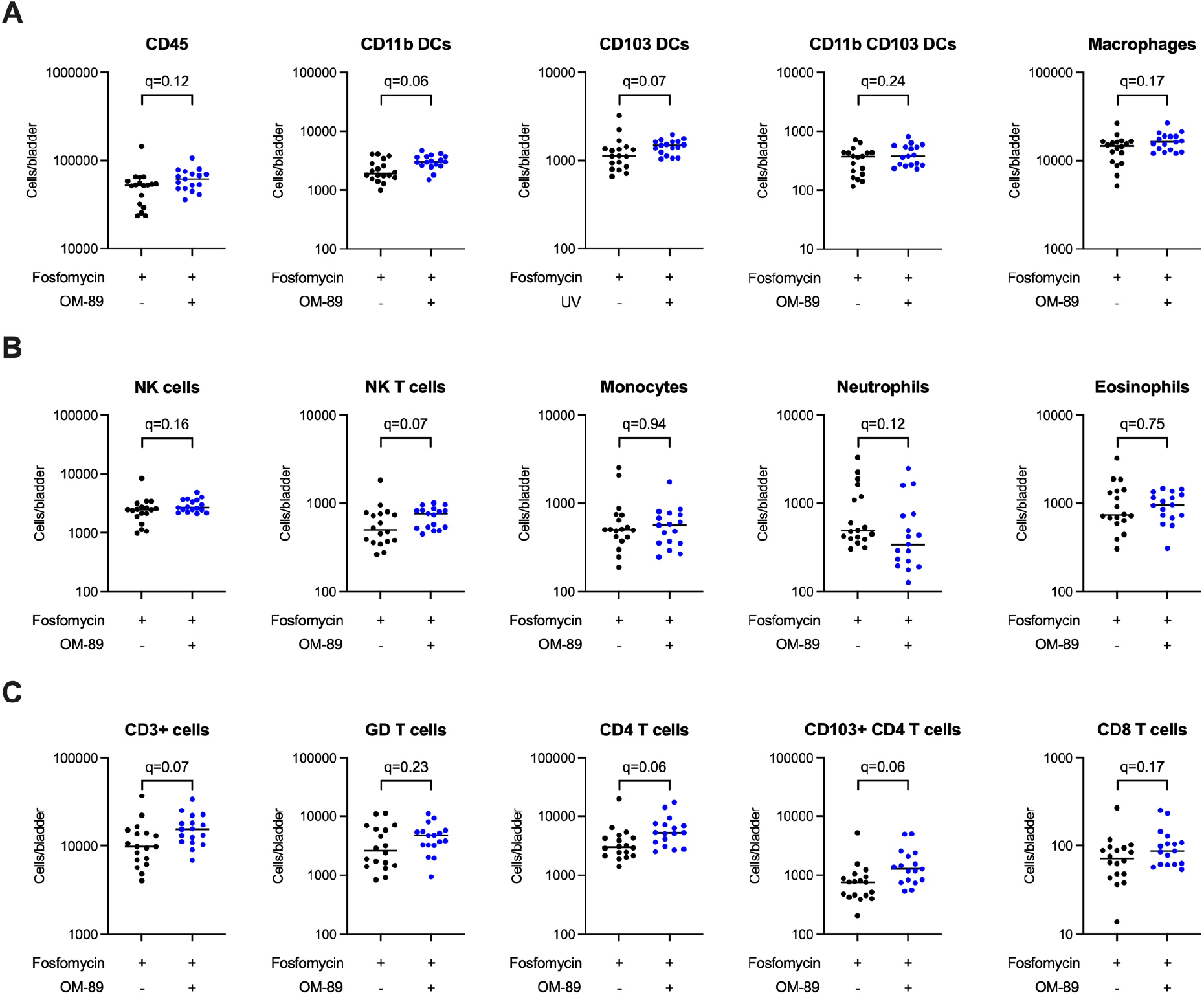
OM-89 treatment does not affect immune cell infiltration in the bladder. Female mice were infected, treated, and sampled as in Figure 2A. Graphs depict the number of indicated immune cells in the bladder. Data are pooled from 3 independent experiments, n=6 mice/experiment. Each dot is a mouse and lines are medians. Nonparametric Mann-Whitney tests comparing fosfomycin treatment to fos/OM-89 treatment for each immune cell were performed. P values were corrected for multiple comparisons using the FDR method and all q values are shown.

### OM-89 does not change effector and memory T cell subsets in the bladder or lymph nodes

We reasoned that although total numbers and proportions of immune cells were not different, fos/OM-89 treatment may have an impact specifically on the development of memory T cell populations. To test this hypothesis, bladders and lymph nodes from 11 day-infected female mice treated with fosfomycin or fos/OM-89 were analyzed by flow cytometry. We observed no significant differences with respect to the total number of CD4 or CD8 T cell subsets present in the bladder (**Figure 5A**) or bladder draining lymph nodes (**Figure 5B**) between the fosfomycin group and the fos/OM-89 group at 11 PI.

**Figure 5:**
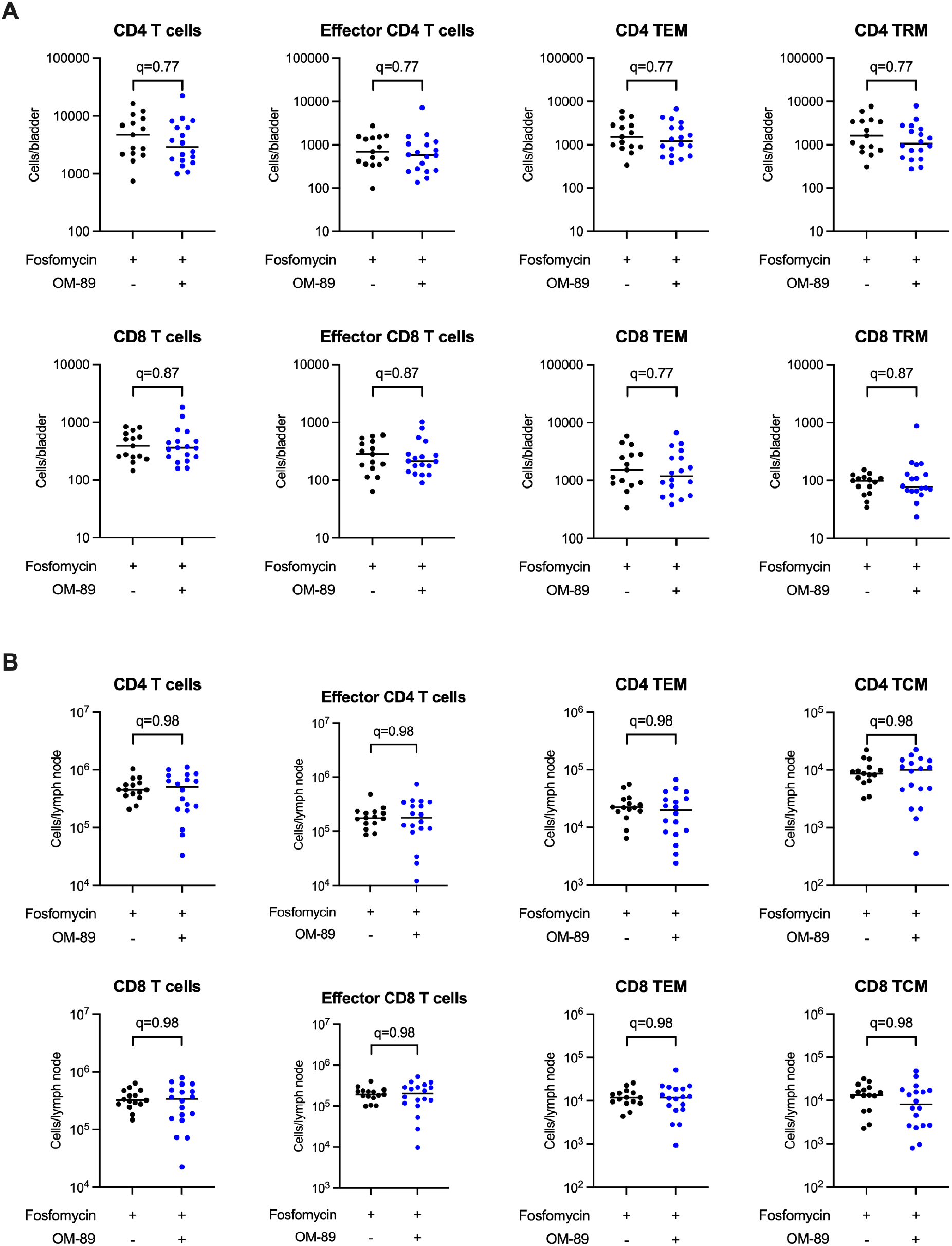
OM-89 treatment does not impact memory and effector T cell subsets in the bladder and bladder draining lymph nodes. Female mice were infected, treated, and sampled as in Figure 2A. Graphs depict the number of indicated immune cells in the (**A**) bladder of (**B**) lymph node. Data are pooled from 3 independent experiments, n=6 mice/experiment. Each dot is a mouse and lines are medians. Nonparametric Mann-Whitney tests comparing fosfomycin treatment to fos/OM-89 treatment for each immune cell were performed. P values were corrected for multiple comparisons using the FDR method and all q values are shown.

### OM-89 treatment does not reduce bacterial reservoirs or improve adaptive immune memory

Finally, we considered that a reduction in bacterial reservoirs may result in fewer recurrence events. Therefore, we infected and treated mice (**Figure 6A**), sacrificing the two groups at 28 days PI. We enumerated the bacterial burden derived from the first infection finding no significant difference in reservoir formation between the fosfomycin treated and the fos/OM-89-treated mice (**Figure 6B**).

**Figure 6:**
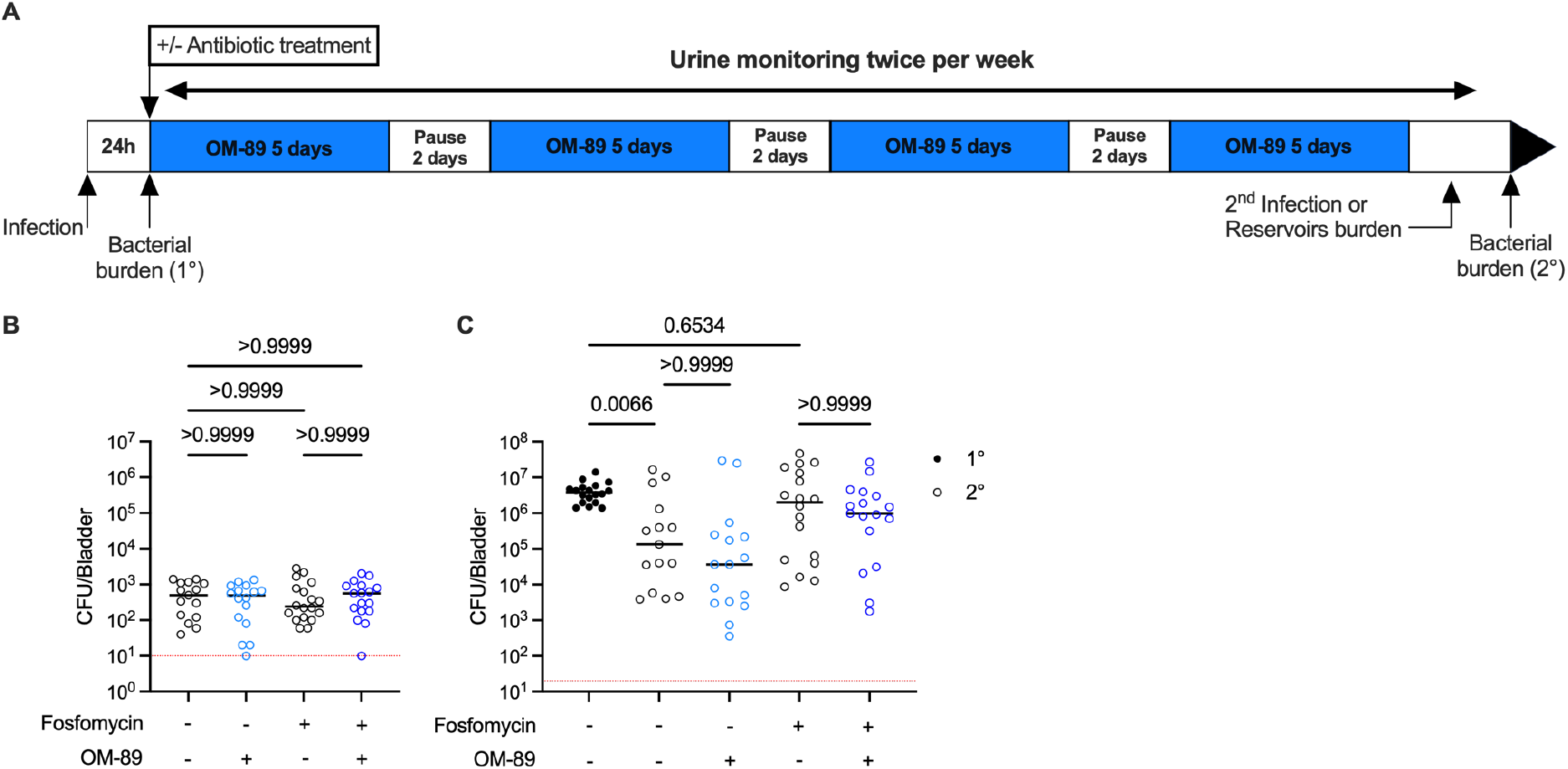
OM-89 does not change bacterial reservoirs or functional immune memory. (**A**) Female mice were infected and then treated with OM-89, fosfomycin, fos/OM-89, or untreaated according to the experimental scheme. (**B-C**) Graphs show bacterial burden in (**B**) reservoirs after 28 days infection or (**C**) 1 day after a second infection. Data are pooled from 3 independent experiments; n=6 mice/experiment. Each dot is a mouse and lines are medians. Red dotted lines display limit of detection. Significance was determined using a Kruskal-Wallis test with a Dunn’s post hoc test to correct for multiple comparisons.

We then considered that the fos/OM-89-treated mice developed immune memory to infection that was not apparent from our immune cell analyses. To test whether fos-OM-89 treated mice were better protected against an intentional recurrent UTI compared to fosfomycin-treated mice, we infected mice, treated them, sampled urine over time, and reinfected the animals at 28 days PI (**Figure 6A**). We previously reported that mice develop a tissue resident memory T cell response that leads to reduced bacterial burden after a second UTI (Rousseau et al., 2023). Here, as expected, bacterial burden was reduced in control, untreated mice after the second infection relative to mice with a primary UTI in the control setting (**Figure 6C**). Interestingly, we observed that fosfomycin treatment abrogated development of immune memory, and bacterial burden was not different between the first and second UTI (**Figure 6C**). This result was similar to our previously reported observations that TMP-SMX treatment also abrogates immune memory to a second UTI by disrupting memory T cell development. Finally, the bacterial burden after a second UTI was not different between OM-89-treated mice and untreated mice or fosfomycin-treated mice and the fos/OM-89 group (**Figure 6C**).

## Discussion

Recurrent urinary tract infection is a major health challenge that severely impacts the quality of life of patients. Recurrent UTI is defined as having an infection twice in six months or three times in 12 months (Kranz et al., 2024). Approximately 25% of all women will experience recurrent UTI during their lifetime (Foxman, 2010) and a vast majority of these infections will be treated with antibiotics, such as TMP-SMX or fosfomycin. The global dissemination of antimicrobial resistance in uropathogens and lack of new antibiotics severely threatens treatment options for UTI and recurrent UTI. As such, alternatives to antibiotics or combination therapies that augment antibiotic efficiency are urgently needed.

OM-89 is thought to act as an immunomodulatory drug, however, its mechanism of action in preventing recurrent UTI is unknown and poorly studied. After administration in Balb/c mice, OM-89 induces serum IgG and IgA responses against the *E. coli* strains used for the preparation of the product (Huber et al., 2000a; Huber et al., 2000b). This is in contrast to our own results, in which we did not observe significant induction of IgG or IgA in the serum or bladder of OM-89-treated mice. The mouse strain may impact the subsequent immune response, particularly as Balb/c mice are considered to be able to mount more prominent antibody responses, generally. Additionally, we measured immunoglobulins within the first 2 weeks of OM-89 treatment, whereas IgA and IgG were measured after 306 days in the previously published study, adding evidence that longer treatment may be necessary to observe adaptive immune changes. In our mouse model, we found that OM-89 treatment, when used in combination with trimethoprim/sulfamethoxypyridazine or fosfomycin, significantly reduced spontaneous recurrent UTI in female mice. In this model, recurrence was defined as presence of bacteria in the urine sample of a mouse with previously sterile urine. Of note, in humans, diagnosis of recurrent UTI includes reporting of symptoms, which is not possible in mice. The strongest effect was observed in combination with fosfomycin in which no spontaneous recurrences were detected. Importantly, OM-89 treatment alone did not reduce spontaneous recurrence events in female mice. These observations support the testing of OM-89 in female patients in combination with antibiotics for the prevention of recurrent UTI. Clinical studies could be designed to identify the best performative OM-89-antibiotic combination for maximum efficiency in reducing recurrent UTI in humans.

Although we identified that OM-89 prevented spontaneous recurrent UTI when used in combination with antibiotics, we did not detect a difference on antibody levels in serum or bladder tissue, circulating immune cell numbers, bladder-infiltrating immune cell numbers, or more specifically on T cell populations known to mediate immunity. We also did not detect a difference in the development of reservoirs or functional immune memory in mice treated with the fosfomycin OM-89 combination compared to OM-89 treated mice. Animals that received OM-89 only were protected to a similar extent as untreated mice, supporting that OM-89 does not prevent spontaneous recurrence by strengthening classical immune memory. Interestingly, we did observe that treatment with fosfomycin abrogated protective memory, as we described for TMP-SMX (Rousseau et al., 2023). Thus, at least two common antibiotics used to treat UTI have a deleterious impact on the memory response to UTI in a pre-clinical model.

OM-89 treatment did not induce detectable changes in innate immune cell infiltration; however it may have induced changes in activation state or responsiveness to infection in these cells. Whether OM-89 impacts other aspects of the innate immune response is unknown. Indeed, OM-89 may have a direct impact on urothelial cells. Urothelial cells detect pathogen-associated molecular patterns (PAMPs) and respond to these stimuli when uropathogens are present in the bladder or when they have invaded the urothelial cell layer (Lacerda Mariano and Ingersoll, 2020). OM-89 treatment may induce production of antimicrobial peptides or upregulate specific molecules to maintain an “alert mode”, which in turn may induce a faster or stronger innate immune response. Thus, direct action of OM-89 on the urothelium is a potentially interesting avenue for future exploration.

In summary, we demonstrated that, in a preclinical mouse of spontaneous recurrent UTI, OM-89 in combination with antibiotics strongly decreased spontaneous recurrences in female mice. Our results support that this is a viable preclinical model, reflecting human disease, to explore the impact of OM-89, and other therapies for the treat of recurrent UTI. This finding will guide current recommendations regarding the target population for this product and clinical protocols to test OM-89 in conjunction with antibiotics to prevent recurrent UTI.

## Methods

### Mice

*In vivo* animal experiments conducted at Institut Pasteur were in accordance with approval of protocol number 2016-0010 by the Comité d’éthique en expérimentation animale Paris Centre et Sud (the ethics committee for animal experimentation), in application of the European Directive 2010/63 EU. Experiments performed at Institut Cochin were approved by the Comité d’Éthique en matière d’Expérimentation Animale Paris Descartes (CEEA 34) under the protocol number APAFiS #34290. C57BL/6J mice, aged 6 weeks were from Charles River Laboratories France. Animals can die during an experiment, potentially due to repeated anesthesia or other factors. In all experiments, n= 6, and when an animal died unexpectedly, its data were removed from the entire experiment.

### Study specific reagents and treatment schedule

A liquid formulation of OM-89 at a concentration of 29.7 mg/mL was stored at +4°C upon reception and for the duration of the treatment phase. Distilled water was used as a vehicle. At the start of each experiment, the OM-89 stock was gently agitated 1 hour and aliquoted and stored at +4°C. Each morning of experimentation, 500 µL of distilled water was added to one Eppendorf tube containing 500 µL of OM-89 for a final concentration of 14.85 mg/mL and gently agitated a minimum of 30 minutes at room temperature prior to administering treatment to the mice. Female and male mice were weighed prior to the start of the experiment and the mean weight of each group was used to calculate the dilution necessary to administer 36 mg/kg OM-89 per mouse. Mice in all groups were weighed once per week for the duration of the experiment. Treatment volumes were recalculated each week based on weight changes in the groups. OM-89 or water were administered *per os* using a P200 pipet and sterile filtered tips, which were changed between each group. Mice were treated with trimethoprim-sulfamethoxypyridazine in the drinking water for 5 days (4 g in 500 mL sterile water) or given a single dose of fosfomycin 200 mg/mL (500 mg/kg) under the tongue using a P200 pipet and sterile filtered tips. Untreated mice received water and water, OM-89, antibiotics, or antibiotic+OM-89 combinations were administered according to the schedules shown in Figure 1A, 2A, or 6A.

### Urinary tract infection and determination of bacterial burden

Female mice were anesthetized by intraperitoneal injection of ketamine (100 mg/kg) and xylazine (5 mg/kg), catheterized transurethrally, and infected with 10^7^ colony forming units (CFU) of uropathogenic *E. coli* (UPEC) strain UTI89-GFP-ampR or UTI89-RFP-kanR in 50 μL PBS (Mora-Bau et al., 2015). Challenge infection was performed 28 days after a primary infection using an isogenic UPEC strain as previously described (Mora-Bau et al., 2015; Zychlinsky Scharff et al., 2019). Resolution was monitored by the presence of bacterial growth from urine samples. Urine was collected every 2-5 days and 2 μL were diluted directly into 8 μL phosphate buffered saline (PBS) spotted on agar plates containing the appropriate antibiotics for the infecting strain. The presence of any bacterial growth was counted as positive for infection. The limit of detection (LOD) for this assay is 500 bacteria per mL of urine. Mice were sacrificed at indicated timepoints, bladders homogenized in sterile PBS, serially diluted, and plated to determine CFU. The LOD for CFU is 20 or 10 per organ for the assessment of acute infection or bacterial reservoirs, respectively.

### Flow cytometry

Bladders were collected at day 11 PI, and prepared for flow cytometry as previously described (Mora-Bau et al., 2015; Rousseau et al., 2023; Zychlinsky Scharff et al., 2019). Single cell suspensions were stained with antibodies shown in Table or S3. Blood was collected in Eppendorf tubes containing 10 µL of EDTA at 100 mM. 10 µL of blood were added to 200 µL of 1step/fix lyse 1X (ebioscience) and 20 µL AccuCheck counting beads (Invitrogen) were added to the same tube to determine absolute cell counts per uL. The remainder of the blood was lysed (BD PharmLyse), resuspended in 100 µL of FACS buffer containing 0.5% of antibodies shown in Table S4. All samples were acquired on a BD LSR Fortessa (BD Biosciences) and analyzed using FlowJo version 10 software.

### Immunoglobulin analysis

Blood was collected at days 0, 5, and 11 PI from the submandibular vein and incubated at room temperature for at least 1 hour. Clotted blood was centrifuged 15 minutes at 1500g 4°C, serum collected and frozen at -20°C in low protein binding 96-well plates until analysis. Bladders were removed at day 11 PI, homogenized, clarified by centrifugation, and stored at -20°C in low protein binding 96-well plates until analysis. After thawing, prior to analysis, samples were centrifuged at 1200 rpm, 4°C, 5 minutes to remove remaining cell debris. All samples were assessed together to avoid inter-assay variability by Millipore MILLIPLEX MAP Mouse Immunoglobulin Isotyping Magnetic Bead Panel -Isotyping Multiplex Assay, according to the manufacturer’s recommendations (Merck Millipore).

### Statistics

Statistical analysis was performed in Prism 10 for Mac OS X using the nonparametric Kruskal-Wallis test with a Dunn’s multiple comparison post-test, or multiple Mann-Whitney tests followed by FDR testing to correct for multiple testing for all data. A p value of less than 0.05 was considered significant. All comparisons made are shown and all corrected *p*-values (*q*-values) are indicated.

## Author contributions

CP, EB, MAI, and MRousseau conceived the study. TC, MD and MRousseau performed experiments and data analysis. MAI and MRousseau wrote the manuscript. TC, MD, EB, CP, RC, MAI, MRousseau reviewed and edited the manuscript. MAI obtained funding and MAI and MRousseau supervised the study. All authors reviewed and approved the manuscript prior to submission.

---

**Table S2:** Antibodies used to analyze bladder immune cells (Figure 4).

| Antibody | Clone | Source |
| --- | --- | --- |
| CD8α | 53-6.7 | BD pharmingen |
| NK1.1 | PK136 | BD pharmingen |
| MHCII | MS/114.15.2 | Invitrogen |
| F4/80 | Cl:A3-1 | Biorad |
| GR1 | RB6-8C5 | BD pharmingen |
| CD11b | M1/70 | BD Horizon |
| CD103 | M290 | BD Horizon |
| CD4 | RM4-4 | Biolegend |
| CD64 | X54-5/7.1 | Biolegend |
| SiglecF | E50-2440 | BD optibuild |
| CD45 | 30-F11 | BD Horizon |
| $\gamma\delta$ TCR | ebioGL3 | Invitrogen |
| CD3 | 145-2C11 | BD Horizon |
| CD11c | HL3 | BD pharmingen |

**Table S3:** antibodies used to analyze T cells subsets in bladder and bladder draining lymph nodes (Figure 5)

| Antibody | Clone | Source |
| --- | --- | --- |
| NK1.1 | PK136 | BD Biosciences |
| CD127 | A7R34 | Invitrogen |
| CD69 | H1.2F3 | BD Biosciences |
| CD8α | 53-6.7 | BD Biosciences |
| Ly6C | HK1.4 | Biolegend |
| **CD103** | M290 | BD Biosciences |
| **CD4** | RM4-5 | Biolegend |
| **CCR7** | 4B12 | BD Biosciences |
| **KLRG1** | 2F1 | BD Biosciences |
| **CD45** | 30-F11 | BD Biosciences |
| **CD44** | IM7 | BD Biosciences |
| **$\gamma\delta$ TCR** | eBioGL3 (GL3) | Invitrogen |
| **CD3** | 145-2C11 | Biolegend |
| **CD62L** | MEL-14 | Invitrogen |

**Table S4:** Antibodies used to analyze blood immune cells (Figure 3).

| **Antibody** | **Clone** | **Source** |
| --- | --- | --- |
| **NK1.1** | PK136 | BD pharmingen |
| **B220** | RA3-6B2 | BD pharmingen |
| **MHCII** | MS/114.15.2 | Invitrogen |
| **CD8α** | 53-6.7 | BD pharmingen |
| **GR1** | RB6-8C5 | BD pharmingen |
| **CD3** | 500A2 | BD pharmingen |
| **CD45** | 30-F11 | BD Horizon |
| **CD115** | AF598 | Invitrogen |
| **SiglecF** | E50-2440 | BD Horizon |
| **CD4** | GK1.5 | ebioscience |

## References

Amoura, A., C. Pistien, C. Chaligne, S. Dion, M. Magnan, A. Bridier-Nahmias, A. Baron, F. Chau, E. Bourgogne, M. Le, E. Denamur, M.A. Ingersoll, B. Fantin, A. Lefort, and I. El Meouche. 2024. Variability in cell division among anatomical sites shapes Escherichia coli antibiotic survival in a urinary tract infection mouse model. Cell Host Microbe 32:900–912 e904.

Anderson, G.G., S.M. Martin, and S.J. Hultgren. 2004. Host subversion by formation of intracellular bacterial communities in the urinary tract. Microbes Infect 6:1094–1101.

Bauer, H.W., S. Alloussi, G. Egger, H.M. Blumlein, G. Cozma, C.C. Schulman, and U.T.I.S.G. Multicenter. 2005. A long-term, multicenter, double-blind study of an Escherichia coli extract (OM-89) in female patients with recurrent urinary tract infections. Eur Urol 47:542–548; discussion 548.

Bauer, H.W., V.W. Rahlfs, P.A. Lauener, and G.S.S. Bleßmann. 2002. Prevention of recurrent urinary tract infections with immuno-active E. coli fractions: a meta-analysis of five placebo-controlled double-blind studies. International Journal of Antimicrobial Agents 19:451–456.

De Nisco, N.J., M. Neugent, J. Mull, L. Chen, A. Kuprasertkul, M. de Souza Santos, K.L. Palmer, P. Zimmern, and K. Orth. 2019. Direct Detection of Tissue-Resident Bacteria and Chronic Inflammation in the Bladder Wall of Postmenopausal Women with Recurrent Urinary Tract Infection. J Mol Biol 431:4368–4379.

Foxman, B. 2010. The epidemiology of urinary tract infection. Nat Rev Urol 7:653–660.

Gadhvi, J.G., P.R.M. Kenee, K.C. Lutz, F. Khan, Q. Li, P.E. Zimmern, and N.J. De Nisco. 2024. Bladder-resident bacteria associated with increased risk of recurrence after electrofulguration in women with antibiotic-recalcitrant urinary tract infection. medRxiv

Huber, M., W. Baier, A. Serr, and W.G. Bessler. 2000a. Immunogenicity of an E. coli extract after oral or intraperitoneal administration: induction of antibodies against pathogenic bacterial strains. Int J Immunopharmacol 22:57–68.

Huber, M., K. Krauter, G. Winkelmann, H.W. Bauer, V.W. Rahlfs, P.A. Lauener, G.S. Blessmann, and W.G. Bessler. 2000b. Immunostimulation by bacterial components: II. Efficacy studies and meta-analysis of the bacterial extract OM-89. Int J Immunopharmacol 22:1103–1111.

Ikaheimo, R., A. Siitonen, T. Heiskanen, U. Karkkainen, P. Kuosmanen, P. Lipponen, and P.H. Makela. 1996. Recurrence of urinary tract infection in a primary care setting: analysis of a 1-year follow-up of 179 women. Clin Infect Dis 22:91–99.

Kranz, J., R. Bartoletti, F. Bruyere, T. Cai, S. Geerlings, B. Koves, S. Schubert, A. Pilatz, R. Veeratterapillay, F.M.E. Wagenlehner, K. Bausch, W. Devlies, J. Horvath, L. Leitner, G. Mantica, T. Mezei, E.J. Smith, and G. Bonkat. 2024. European Association of Urology Guidelines on Urological Infections: Summary of the 2024 Guidelines. Eur Urol 86:27–41.

Lacerda Mariano, L., and M.A. Ingersoll. 2020. The immune response to infection in the bladder. Nat Rev Urol 17:439–458.

Lacerda Mariano, L., M. Rousseau, H. Varet, R. Legendre, R. Gentek, J. Saenz Coronilla, M. Bajenoff, E. Gomez Perdiguero, and M.A. Ingersoll. 2020. Functionally distinct resident macrophage subsets differentially shape responses to infection in the bladder. Sci Adv 6:

Mora-Bau, G., A.M. Platt, N. van Rooijen, G.J. Randolph, M.L. Albert, and M.A. Ingersoll. 2015. Macrophages Subvert Adaptive Immunity to Urinary Tract Infection. PLoS Pathog 11:e1005044.

Mysorekar, I.U., and S.J. Hultgren. 2006. Mechanisms of uropathogenic Escherichia coli persistence and eradication from the urinary tract. Proc Natl Acad Sci U S A 103:14170–14175.

Prattley, S., R. Geraghty, M. Moore, and B.K. Somani. 2020. Role of Vaccines for Recurrent Urinary Tract Infections: A Systematic Review. Eur Urol Focus 6:593–604.

Rosen, D.A., T.M. Hooton, W.E. Stamm, P.A. Humphrey, and S.J. Hultgren. 2007. Detection of intracellular bacterial communities in human urinary tract infection. PLoS Med 4:e329.

Rosen, D.A., J.S. Pinkner, J.N. Walker, J.S. Elam, J.M. Jones, and S.J. Hultgren. 2008. Molecular variations in Klebsiella pneumoniae and Escherichia coli FimH affect function and pathogenesis in the urinary tract. Infect Immun 76:3346–3356.

Rousseau, M., L. Lacerda Mariano, T. Canton, and M.A. Ingersoll. 2023. Tissue-resident memory T cells mediate mucosal immunity to recurrent urinary tract infection. Sci Immunol 8:eabn4332.

Schilling, J.D., R.G. Lorenz, and S.J. Hultgren. 2002. Effect of trimethoprim-sulfamethoxazole on recurrent bacteriuria and bacterial persistence in mice infected with uropathogenic Escherichia coli. Infect Immun 70:7042–7049.

Worby, C.J., H.L.t. Schreiber, T.J. Straub, L.R. van Dijk, R.A. Bronson, B.S. Olson, J.S. Pinkner, C.L.P. Obernuefemann, V.L. Munoz, A.E. Paharik, P.N. Azimzadeh, B.J. Walker, C.A. Desjardins, W.C. Chou, K. Bergeron, S.B. Chapman, A. Klim, A.L. Manson, T.J. Hannan, T.M. Hooton, A.L. Kau, H.H. Lai, K.W. Dodson, S.J. Hultgren, and A.M. Earl. 2022. Longitudinal multiomics analyses link gut microbiome dysbiosis with recurrent urinary tract infections in women. Nat Microbiol 7:630–639.

Yamamoto, S., T. Tsukamoto, A. Terai, H. Kurazono, Y. Takeda, and O. Yoshida. 1997. Genetic evidence supporting the fecal-perineal-urethral hypothesis in cystitis caused by Escherichia coli. J Urol 157:1127–1129.

Yang, X., H. Chen, Y. Zheng, S. Qu, H. Wang, and F. Yi. 2022. Disease burden and long-term trends of urinary tract infections: A worldwide report. Front Public Health 10:888205.

Zychlinsky Scharff, A., M.L. Albert, and M.A. Ingersoll. 2017. Urinary Tract Infection in a Small Animal Model: Transurethral Catheterization of Male and Female Mice. J Vis Exp

Zychlinsky Scharff, A., M. Rousseau, L. Lacerda Mariano, T. Canton, C.R. Consiglio, M.L. Albert, M. Fontes, D. Duffy, and M.A. Ingersoll. 2019. Sex differences in IL-17 contribute to chronicity in male versus female urinary tract infection. JCI Insight 5:

