## Supplementary figure 1 for "Oral OM-89 in combination with antibiotics prevents recurrent infection in a mouse model of urinary tract infection"

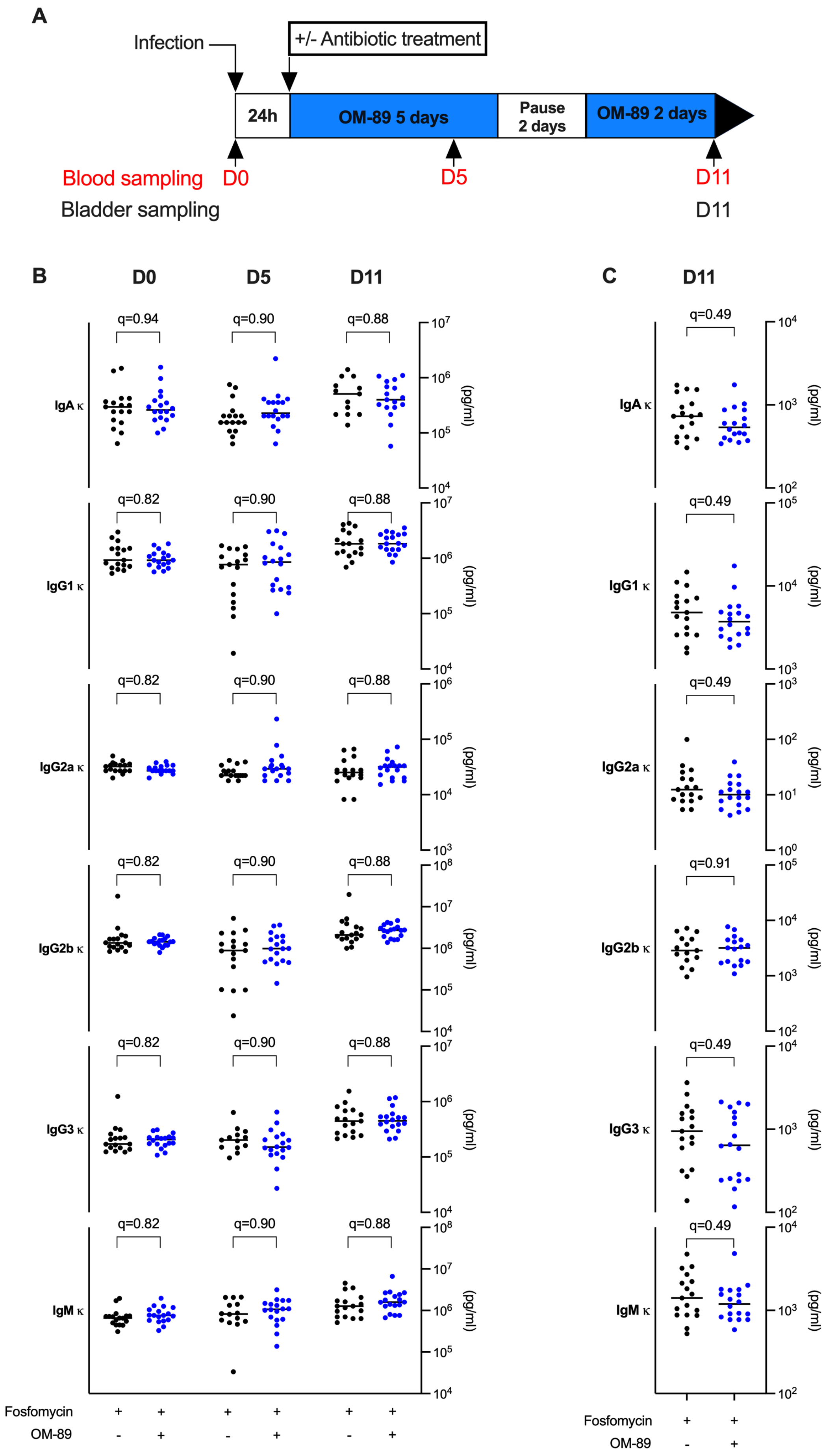
A
Infection
+/- Antibiotic treatment
24h
OM-89 5 days
Pause 2 days
OM-89 2 days
Blood sampling D0
D5
D11
Bladder sampling
D11
B
D0
D5
D11
IgA κ
IgG1 κ
IgG2a κ
IgG2b κ
IgG3 κ
IgM κ
q=0.94
q=0.90
q=0.88
q=0.82
(pg/ml)
Fosfomycin + + + + + +
OM-89 - + - + - +
C
D11
q=0.49
q=0.91
(pg/ml)
Fosfomycin + +
OM-89 - +

**Supplementary Figure 1: OM-89 does not alter antibody levels in serum or bladder. (A)** Female mice were infected and then treated with fosfomycin or fos/OM-89 according to the experimental scheme. Graphs show antibody concentrations in (**B**) serum on day 0, 5, and 11 PI and (**C**) bladder homogenates on day 11 PI. Data are pooled from 3 independent experiments; n= 6 mice/experiment. Each dot is a mouse and lines are medians. Nonparametric Mann-Whitney tests comparing fosfomycin treatment to fos/OM-89 treatment for each immunoglobulin concentration were performed. P values were corrected for multiple comparisons using the FDR method and all q values are shown.
